# Mobilization of Pack-CACTA transposons in Arabidopsis reveals the mechanism of gene shuffling

**DOI:** 10.1101/342816

**Authors:** Marco Catoni, Thomas Jonesman, Elisa Cerruti, Jerzy Paszkowski

## Abstract

Pack-TYPE transposons are a unique class of potentially mobile non-autonomous elements that can capture, merge and relocate fragments of chromosomal DNA. It has been postulated that their activity accelerates the evolution of host genes. However, this important presumption is based only on the sequences of currently inactive Pack-TYPE transposons and the acquisition of chromosomal DNA has not been recorded in real time. We have now for the first time witnessed the mobilization of novel Pack-TYPE elements related to the CACTA transposon family over several plant generations. Remarkably, these elements tend to insert into genes as closely spaced direct repeats and they frequently undergo incomplete excisions, resulting in the deletion of one of the end sequences. These properties constitute a mechanism of efficient acquisition of genic DNA residing between neighbouring Pack-TYPE transposons and its subsequent mobilization. Our work documents crucial steps in the formation *in vivo* of novel Pack-TYPE transposons and thus the mechanism of gene shuffling mediated by this type of mobile element.

## Introduction

Autonomous transposable elements (TEs) encode all factors required for transposition, whereas non-autonomous elements react to factors provided *in trans* (Lisch, 2013). In DNA transposons, terminal inverted-repeat sequences (TIRs) of transposition-competent non-autonomous TEs are recognized by a transposase of a related autonomous element (Skipper et al., 2013). The extremities of Pack-TYPE elements include TIRs, while their internal sequences are derived from host chromosomes (Bennetzen, 2005; Jiang et al., 2011; Kawasaki and Nitasaka, 2004; Yu et al., 2000; Zabala and Vodkin, 2005). For example, Pack-MULEs are Pack-TYPE elements that populate the rice genome (Jiang et al., 2004), with approximately 3000 Pack-MULEs carrying segments of about 1000 different genes (Jiang et al., 2011). This heterogeneous population of transposons has had a major impact on the current organization of rice chromosomes and the evolution of rice genes (Jiang et al., 2004). Structures resembling Pack-TYPE transposons are also present in Arabidopsis (Gilly et al., 2014; Yu et al., 2000), where epigenetic suppression mediated by DNA methylation appears to render them immobile (Lister et al., 2008). Elimination of DNA METHYLTRASFERASE1 (MET1) activates the genome-wide transcription of TEs by erasing CpG methylation (Zilberman et al., 2007). Importantly, the absence of CpG methylation persists over many plant generations, even after restoration of MET1 activity (Catoni et al., 2017; Kankel et al., 2003; Saze et al., 2003). This observation prompted the construction of epigenetic recombinant inbred lines (epiRILs) from a cross between two isogenic parents: the wild type and a *met1* mutant deficient in CpG methylation (Reinders et al., 2009). F2 progeny with MET1 activity were further propagated by single-seed descent for eight generations, resulting in 68 epiRILs with mosaic methylation patterns (Reinders et al., 2009). Here, we screened for new TE mobilized in epiRILs, and we identified a new family of Pack-TYPE mobile elements with CACTA1-derived Terminal Inverted Repeats (TIRs). The study of real time mobilization of one of these new Pack-CACTA elements allowed proposing the mechanism of gene shuffling by this type of transposon.

## Results

DNAseq of PCR-free genomic libraries for the 68 epiRILs was used to search for TE mobilization, which is usually associated with increased copy number (Chuong et al., 2017; Lisch, 2013). We looked for TE-annotated loci for which read coverage was at least 1.8-fold higher than in wild-type Col-0. We then compared putative activity levels of these TEs by the average increase in sequencing coverage. According to this analysis, retrotransposon EVADE (EVD) (Mirouze et al., 2009) was the most active, with an average 11.8-fold coverage increase (Supplementary Table 2). Surprisingly, the second most active element, with a 6.2-fold increase, had an unusually complex structure of two short terminal DNA stretches of 451 bp (AT4TE18505) and 337 bp (AT4TE18510) (annotated as a member of the ATENSPM3 family of DNA transposons) separated by a sequence annotated as gene AT4G07526 of unknown function (Figure 1A). The putative product of this gene showed no similarity to any known transposase and the overall transposon structure, residing on chromosome 4, resembled a non-autonomous Pack-TYPE element.

**Figure 1.**
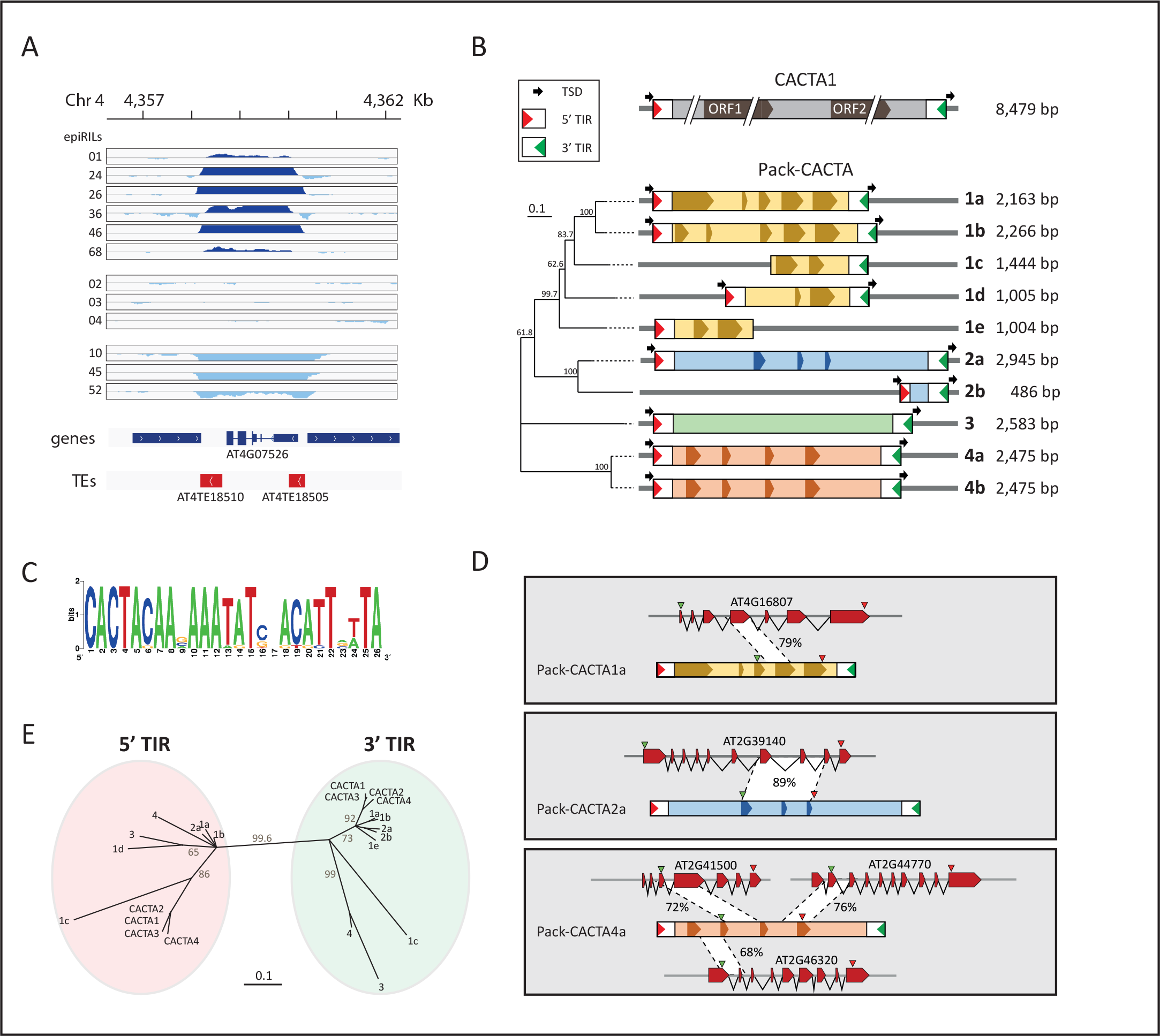
A novel non-autonomous pack-TYPE transposon related to CACTA becomes mobile in epiRILs. A) Copy number changes in epiRILs of a Pack-TYPE element (Pack-CACTA1a) residing on chromosome 4. EpiRIL numbers are indicated left of each track. Differences to reference Col-0 control in log-transformed coverage of reads plotted in the interval of −1 to 1 are mapped to the Pack-CACTA1a for each track. TAIR10 annotation of genes (blue) and TEs (red) is displayed at the bottom. B) Structure of Pack-CACTA family. Four groups of elements are marked in different colours with exons represented by a darker hue. The phylogenetic tree was obtained using alignment of full-length sequences. Numbers at each node indicate bootstrap support values of 1000 replications. The 5’ terminal inverted repeat (5’ TIR) is marked as a red triangle and 3’ TIR as a green triangle. Target site duplications of 3 bp (TSD) are marked as black arrows. C) Sequence logo (http://weblogo.berkeley.edu/logo.cgi) obtained for alignment of TIRs of the 10 members of the Pack-CACTA family. D) Chromosomal origin of Pack-CACTA sequences. The percentage identities to Arabidopsis genes are indicated for each area. E) Sequence relationships of the TIRs of Pack-CACTA groups. CACTA1 is annotated as AT2TE20205, CACTA2 as AT1TE42210, CACTA3 as AT2TE18415, and CACTA4 as AT1TE36570. Numbers at each node indicate bootstrap support values of 1000 replications.

AT4TE18505 and AT4TE18510 contain TIRs beginning with 8 base pairs (CACTACAA) that are also a feature of the CACTA1 autonomous transposon (Miura et al., 2001), which is active in the epiRILs (Reinders et al., 2009) (Supplementary Table 2). A blast search with 150 bp of both terminal sequences identified nine additional elements in the Col-0 genome with similar structures and CACTA1-like termini (Figure 1B). These included conserved terminal sequences of 26 bp (Figure 1C) and a 3-bp target site duplication (TSD) (Supplementary Table 3). Their 150-bp terminal sequence identities ranged from 62 to 95.3% (Supplementary Table 4). Five of the elements shared related DNA sequences located between the CACTA1-like termini, while the remaining five elements carried sequences of different chromosomal origin (Figure 1B and D). Interestingly, not all of the captured sequences displayed similarities to Arabidopsis genes (Figure 1D) and DNA sequence databases did not reveal their origins. The TIR sub-terminal sequences of CACTA family transposons differ slightly between the 5’ and the 3’ ends (Frey et al., 1990). These differences were also present in the possibly active Pack-TYPE elements (Figure 1E) we discovered. Therefore, we called this family Pack-CACTA, which in the Arabidopsis Col-0 accession consists of 10 TEs divided up, according to their sequence similarities, into four groups (Figure 1B).

We found 50 new insertions of Pack-CACTA1a and 3 new insertions of Pack-CACTA2a in 8 and 3 epiRILs, respectively (Supplementary Table 5). In line epi26, Pack-CACTA1a and Pack-CACTA2a appeared mobile (Supplementary Table 5). New transposed copies were found mostly within euchromatic chromosomal regions (65%), resembling the insertion preference of CACTA1 (Supplementary Figure 2A). We validated new insertions of Pack-CACTA1a by locus-specific PCR, choosing 11 random sites in epi26 and epi46 and two sites with Pack-CACTA2a insertions in epi14 and epi26 (Supplementary Figure 2B, C and Supplementary Table 6). Thus, both Pack-CACTA1a and Pack-CACTA2a were currently transposing and generating TSD of three nucleotides (Supplementary Data 1).

To better examine the timing and consequences of Pack-CACTA1a transposition, we characterized in more detail five new insertion sites in 10 sibling plants each of epi26 and epi46 (Supplementary Figure 3 A,B, C). We recorded the presence or the absence of Pack-CACTA1a in individual sibling plants (Supplementary Figure 3D, E). The results are consistent with genetic segregation due to the initial hemizygosity of recent insertions and/or the frequent excision of newly inserted transposons (Figure 2A). In the case of transposon excision, we expected footprints that would be absent in the corresponding wild-type locus. In different epi26 sibling plants, we rescued sequences of both intact target loci and those containing footprints of the transposon (Figure 2B and Supplementary Data 2). The results are consistent with high transpositional activity of Pack-CACTA1a in epiRILs during inbreeding as well as somatic excision in tissues of individual plants.

**Figure 2.**
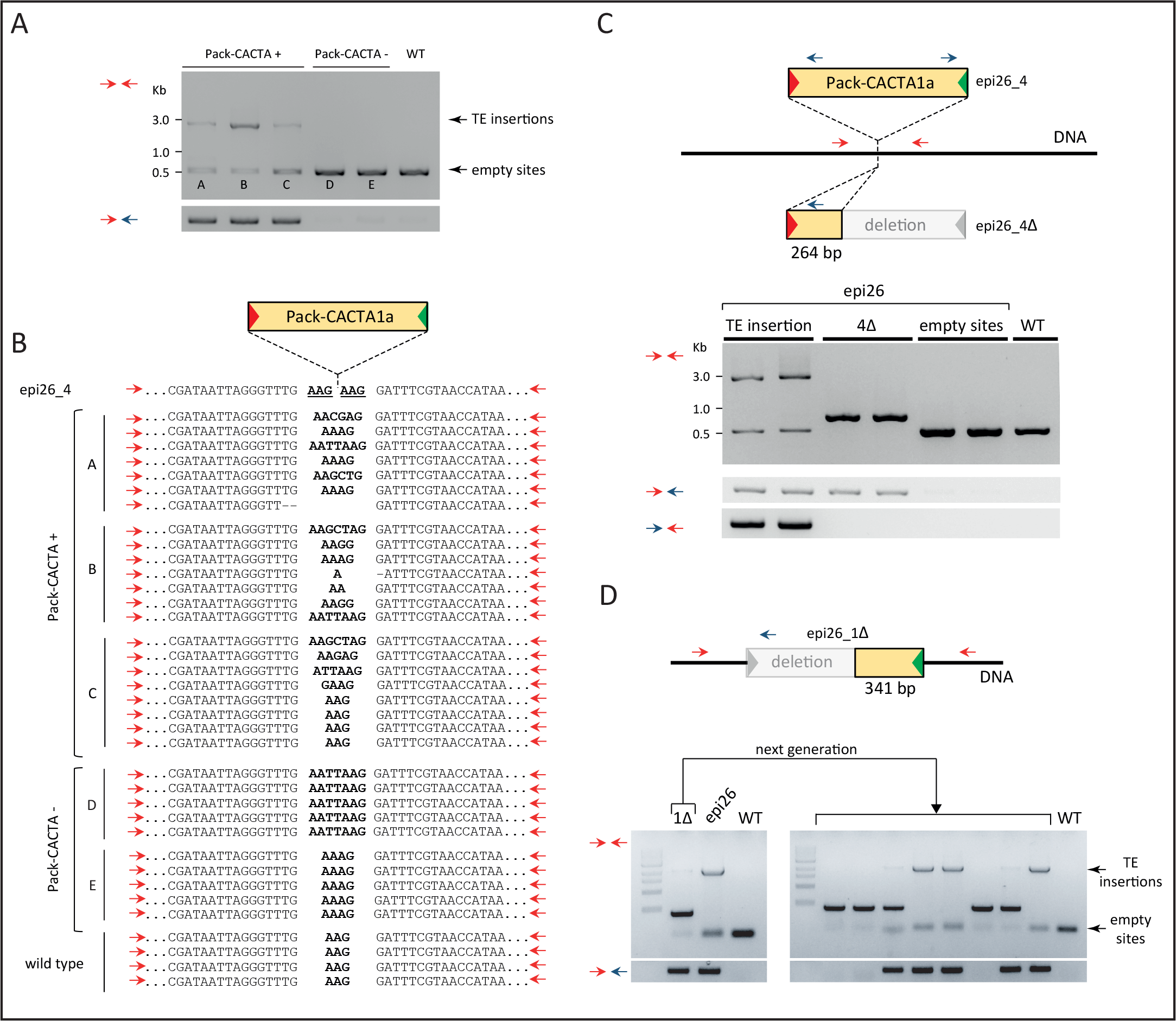
Hallmarks of Pack-CACTA1a excisions. A) Excisions of Pack-CACTA1a from locus 4 of epiRIL26 (epi26_4) in five epi26 plants. Gel-separated PCR products were obtained with primers depicted in panel C as red or blue arrows. Lines marked as “Pack-CACTA +” have a locus 4 containing a Pack-CACTA1a insert (indicated as “TE insertions“) and in plants marked as “Pack-CACTA -“Pack-CACTA1a underwent excision, resulting in “empty sites”. The wild-type Col-0 sample is marked as WT. B) DNA sequences of PCR amplified “empty sites” displayed in panel A. The sequence of the initial epi26_4 containing the TE insertion is provided in the first line and TSD of 3 bp is underlined. Letters A-E left of the sequences correspond to the labelling in panel A. C) Identification of an aberrant excision of Pack-CACTA1a from epi26_4. Two lines of the gel, marked as 4Δ, represent epi26 plants that underwent aberrant excisions resulting in terminal deletion. D) Identification of an additional aberrant excision event of Pack-CACTA1a (left panel) at a different locus of epi26 plants (epi26_1) and its transgenerational inheritance (right panel). All symbols and markings are as in panel C. The line marked 1Δ represents a plant homozygote for a transposon insertion at epi26_1, in which only one Pack-CACTA1a was aberrantly excised. The remaining full-length transposon is not visible due to PCR competition. The ladder is 1 kb NEB.

Unexpectedly, in some plants of epi26 locus 4 (epi26_4) (Supplementary Figure 3F) we found a 264-bp fragment of the CACTA1a 5’ sequence flanked by 3 bp of TSD (Figure 2C and Supplementary Data 3). Ten epi26 plants with a Pack-CACTA1a insertion at this locus contained either the entire Pack-CACTA1a or its deletion derivative (Supplementary Figure 3F). Therefore, it is most probable that the deletion resulted from aberrant excision of a full copy from this location. Importantly, we did not detect somatic “reversions” to a wild type-like locus in plants homozygous for the deleted version of Pack-CACTA1a. The fragmented element was apparently immobilized at this new location (Figure 2C).

To investigate the frequency and features of aberrant excisions of Pack-CACTA1a, we examined a further 64 sibling plants each of epiRILs epi26 and epi46, searching for additional deletion events at five loci at which complete insertions of Pack-CACTA1a were previously found. To this end, we used locus-specific primers that flanked each of the putative Pack-CACTA1a target sites (Supplementary Figure 3C). Six independent deletion events were detected in 640 PCR reactions, two in epi26 and four in epi46 (Figures 2D, Supplementary Figure 4). Therefore, generation of deletion derivatives of this transposon appear to be quite common: in approximately 5% of plants or in 1% of loci previously targeted by Pack-CACTA1a, and at least some of these are transgenerationally transmitted (Figure 2D and Supplementary Figure 4C)

In the case of transposon insertion in an active gene, TIR deletion could stably alter expression of the gene and/or alter the structure of its transcript. Indeed, in one case we detected a novel hybrid mRNA initiated within a promoter residing in the remaining part of Pack-CACTA1a, the second exon of the Pack-CACTA1a “passenger gene” was fused with the exon of a downstream chromosomal gene (Figure 3 left panels and Supplementary Data 3). In a second case, the 3’ part of the transposon was inserted in a gene exon; thus, its sequence formed part of the mature gene transcript (Figure 3 right panels and Supplementary Data 4).

**Figure 3.**
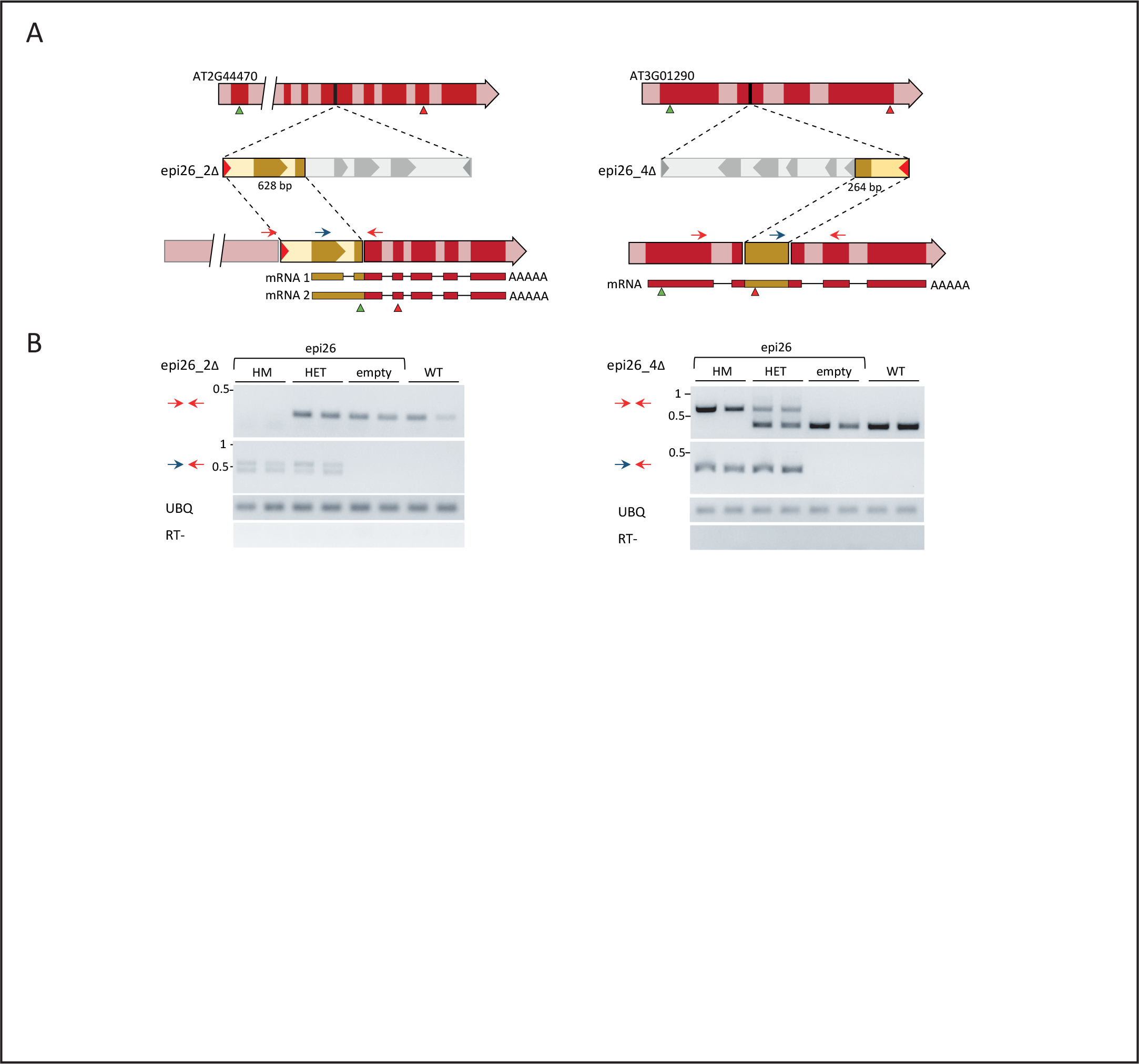
Deletion derivatives of Pack-CACTA1a alter gene transcripts. A) Structures of new Pack-CACTA1a inserts that underwent aberrant excisions. Deleted parts of Pack-CACTA1a are in grey. Exons are in darker and introns in lighter hues. Primers used for RtPCR are shown as blue and red arrows. Deduced, novel mRNAs are displayed at the bottom, with green and red triangles representing initiation and termination codons, respectively. B) Results of RT-PCR performed on single epi26 homozygous (HM), heterozygous (HET) or wild typelike (empty) plants for the corresponding Pack-CACTA1a deletion derivatives depicted in A. The combination of primers used is shown on the left. Col-0 wild-type plants are shown as control (WT). Amplification of ubiquitin mRNA (UBQ) and samples without reverse transcriptase (RT-) were used as controls. Two biological replicates per genotype were used.

The sequences of remaining fragments of the transposon showed that deletions occurred at approximately equal frequencies at the 3’ (3 deletions) and 5’ terminals (4 deletions) (Figure 2C,D, Supplementary Figure 4, Supplementary Data 5). This observation suggested that incorporation of chromosomal DNA into a novel Pack-CACTA element requires the presence of two neighbouring transposons inserted in the direct orientation. If such a pair of Pack-Type transposons, separated by a short stretch of chromosomal DNA, undergoes complementary deletions of each of the elements, a “hybrid” Pack-CACTA encompassing chromosomal DNA within the respective 5’ and 3’ terminals of the former transposons would be generated (Figure 4A). To examine whether Pack-CACTA1a integrates frequently at closely linked chromosomal locations, we designed a PCR-based screen to recover chromosomal DNA stretches residing between putative neighbouring new inserts of Pack-CACTA1a (Figure 4B). Obviously, this approach can only detect insertions separated by relatively short stretches of chromosomal DNA. Therefore, it was surprising that numerous amplifications of chromosomal DNA fragments were observed in the 128 DNA samples of epi26 and epi46 used previously for the Pack-CACTA1a deletion screen (Figure 4C and Supplementary Figure 5). The sequences of 28 different PCR products from 20 plants (Supplementary Figure 6 and Supplementary Data 6, 7) all contained fragments of Arabidopsis genomic DNA of various sizes (38 to 2,137 bp) between two Pack-CACTA1a copies. Remarkably, 27 PCR products revealed insertions of Pack-CACTA1a as tandems in direct orientation and only one inversion (Supplementary figure 6). The frequent formation by newly inserted copies of Pack-CACTA1a of direct repeats interspaced by short stretches of chromosomal DNA, combined with the observed formation of terminal deletion derivatives of the transposon, suggest that pairs of Pack-CACTA1a may form new “hybrid” elements. Indeed, we recovered three deletion derivatives of Pack-CACTA1a among the 27 direct transposon repeats (Supplementary Figure 6). Importantly, most neighbouring Pack-CACTA1a insertions were separated by genic sequences (21 out of 28, Supplementary Figure 6); in one case three Pack-CACTA1a insertions were located in one gene (Figure 4D).

**Figure 4.**
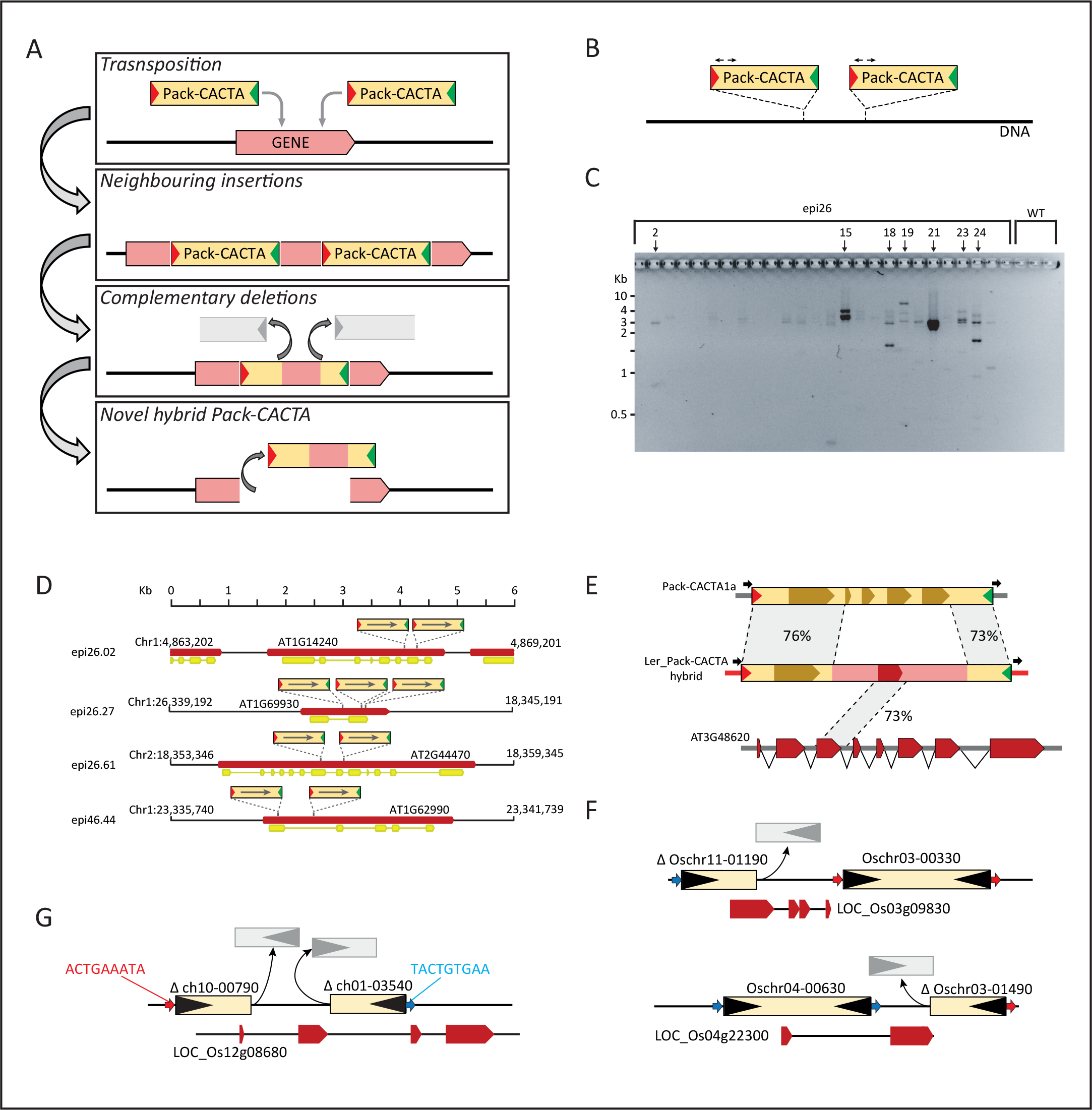
High frequency of Pack-CACTA1a insertion at some loci support a model for DNA capture in new pack-TYPE transposons. A) Possible mechanism of genic DNA capture by Pack-CACTA1a. New transposon insertions form a closely spaced tandem array. Subsequently, neighbouring elements undergo aberrant excisions resulting in complementary, terminal deletions, which form a novel hybrid Pack-CACTA element that incorporates chromosomal DNA which separated the initial closely spaced insertions. B) PCR strategy used for detection of neighbouring insertions of Pack-CACTA1a. Black arrows indicate the primers. C) Results of a PCR screen for neighbouring insertions of Pack-CACTA1a in 30 epi26 plants. Cloned fragments corresponding to adjacent insertions are marked with black arrows. The full set for all tested plants is displayed in Supplementary Figure 5. D) Arrangement of Pack-CACTA1a copies in four representative loci containing tandem insertions (data for other loci are presented in Supplementary Figure 6). The orientation of Pack-CACTA1a copies is indicated with an arrow (5’ to 3’ direction). Red and yellow rectangles represent genes and their transcripts, respectively. E) Structure of a Col-0 Pack-CACTA1a-related element identified in the Ler accession. The percentage sequence identities with Pack-CACTA1a and the acquired gene AT3G48620 are indicated. F) Examples of rice Pack-MULE terminal deletion derivatives neighbouring intact Pack-MULE elements. Black triangles represent TIRs. Blue and red arrows represent 9 bp TSD (different colours correspond to different sequences). Red rectangles represent coding regions of genes. G) Example of neighbouring Pack-MULE deletion derivatives compatible with the formation of a novel hybrid Pack-MULE element. Symbols and colours as in F.

Obvious extrapolations from the observed properties of active Pack-CACTA1a provide crucial clues as to the mechanism by which Pack-TYPE transposons acquire and relocate stretches of chromosomal DNA (Figure 4A). However, detection of a newly formed, transpositionally active Pack-CACTA1a derivative would require the screening of several thousand plants harbouring currently active parental elements.

As an alternative, we searched for structural variants of Pack-CACTA1a elements in the newly assembled genome of the Arabidopsis Ler-0 accession and retrieved eight analogous elements (Supplementary Figure 7 and Supplementary Table 7). Of these, the sequences of three full-length elements and two deletion derivatives were closely related to Pack-CACTA1a of Col-0, with sequence identities of 73 to 90% (Supplementary Table 7). The sequences of the remaining three copies of Ler-0 Pack-CACTA elements resembled Pack-CACTA1a only in the first 992 bp and the last 325 bp with 76% and 72% identities, respectively. Remarkably, the internal 1,181 bp showed no similarity to the “parental” element but contained a 132-bp stretch with 73% identity to the second exon of gene AT3G48620 encoding a putative protein of the outer membrane family (OMP85) (Figure 4E). This novel structure of a Ler-0 relative of Pack-CACTA1a suggests that a Pack-CACTA transposon in this Arabidopsis accession could capture and move chromosomal DNA by a mechanism compatible with that we recorded in real time for Pack-CACTA1a transposition in Col-0.

In rice, previous analyses of Pack-MULE transposons considered only complete elements with both terminal repeats encompassing various fragments of rice chromosomal DNA (Jiang et al., 2004). We reanalysed available rice genomic sequences for intermediates of chromosomal DNA acquisition by rice Pack-MULEs, such as transposon deletions due to aberrant excision events. We applied a blast search across the rice genome using as a query 2,853 previously retrieved pack-MULE intact elements (Jiang et al., 2004) and filtered results to recover incomplete elements with only one terminal sequence (see Methods). Of the 2,151 entries meeting these criteria (Supplementary Data 8), 269 (13 %) Pack-MULE deletion derivatives resided less than 10 kb from an intact Pack-MULE element (Figure 4F). Pack-MULEs consisting of terminal sequences flanked by different target site duplications, making up 17% of the Pack-MULE population, were previously discarded as false positives (Jiang et al., 2004). However, considering the deduced mechanism of DNA acquisition by Pack-TYPE elements, part of these putative transposons may represent cases of newly generated but not yet relocated hybrid Pack-MULEs consisting of terminal sequences from two different parental elements that underwent aberrant excisions (Figure 4G). Thus, the abundant Pack-MULEs of the rice genome, which have contributed significantly to its current organization, seem to include numerous examples of putative transposon intermediates consistent with the scheme of chromosomal DNA acquisition during real-time Pack-CACTA mobilization in Arabidopsis (Figure 4A).

## Discussion

A massive contribution of Pack-TYPE transposon activity to the evolution of plant genes and genomes was deduced from analyses of rice, Brassica and soybean (Alix et al., 2008; Hanada et al., 2009; Jiang et al., 2004; Jiang et al., 2011; Zabala and Vodkin, 2005). It was postulated that repair of nicks and gaps created at the transposon insertions sites or arising due to the element structures might be responsible for the acquisition of chromosomal DNA (Bennetzen and Springer, 1994; Engels et al., 1990; Yamashita et al., 1999). Such DNA-repair mechanisms would operate prior transposon relocation and the newly generated elements would than move to different chromosomal locations only after incorporation of chromosomal DNA. Here, when studying transposition properties of Pack-CACTA1a elements *in vivo*, we found that acquisition of chromosomal DNA seems to be tightly linked to the transposition. Pack-CACTA1a transposition has a marked tendency to produce closely spaced repeats in direct orientation. Subsequently, complementary aberrant excisions of the elements within these repeats, leaving behind one of the two TIRs, generate a novel “hybrid” Pack-CACTA that incorporates the chromosomal DNA that separated the two neighbouring insertions (Figure 4A).

This quite simple and efficient mechanism of chromosomal DNA acquisition relies on the combination of known characteristics of particular DNA transposons. For example, maize Ac/Ds, P-element of *Drosophila*, Tc1 transposon in *Caenorhabditis* and *Sleeping Beauty* in mouse tend to transpose to nearby locations, termed “local hopping” (Bancroft and Dean, 1993; Carlson et al., 2003; Fischer et al., 2003; Skipper et al., 2013; Zhang and Spradling, 1993). However, the orientation of elements in such integration arrays are random and not preferentially direct, as observed for Pack-CACTA1a. Incomplete or aberrant excision is known for CACTA-like transposable elements in soybean (Xu et al., 2010), while maize chromosomal aberrations, like breakages and fusions, were previously associated with the transposition of TIRs belonging to two different neighbouring Ac/Ds elements (Ralston et al., 1989). It is interesting that Pack-CACTA1a combines both properties, i.e. efficient formation of closely linked tandem insertions in direct orientation and frequent aberrant excisions that lead to complementary deletion of one of the two TIRs of each pair. The remaining terminals of the neighbouring transposons encompass a new, single element incorporating the intervening chromosomal DNA.

The incorporation of gene fragments and their duplication by transposition to new genomic positions has been suggested for certain TE families, including MULEs, CACTAs and HELITRONs (Lisch, 2013). This “transduplication” process seems to be very frequent in rice and can be clearly assigned to the historical activity of Pack-MULEs (Juretic et al., 2005). Importantly, transduplicated genes seem to be under purifying selection (Hanada et al., 2009) or serve regulatory functions (Juretic et al., 2005). This implies that Pack-TYPE transposon mobilization may have a direct and profound influence on gene evolution, generating new gene functions or regulatory activities by shuffling parts of various coding regions across the genome. Taking into account the high rates of nearby directional insertion and aberrant excision observed here for the Pack-CACTA1a element, the “TE-assisted” origin of genes may be strongly underestimated, especially in plants with genomes larger than Arabidopsis.

In hypomethylated Arabidopsis inbred lines, we followed for several plant generations mobilization of novel Pack-TYPE elements related to the family of CACTA transposons and we witnessed the initial steps of chromosomal DNA acquisition. Our unique data *in vivo* allowed proposing the mechanism of gene shuffling by this type of transposon. This mechanism is directly linked to their transposition properties and therefore not only depending by DNA repair activities of the host, as it has been postulated so far.

## Methods

### Plant material

The *met1-3* derived epiRIL population was described previously (Reinders et al., 2009). From this population, the 9^th^ inbred generation was grown to produce the 68 epiRILs used in this study. Col-0 and 2^nd^ generation *met1-3* plants (Saze et al., 2003) were used as controls. Unless indicated otherwise, plants were grown in soil under long-day conditions (21°C, 16 h light, 8 h dark).

### DNA extraction and library preparation

Arabidopsis seeds were sown on ½ MS (Plant) medium plates and grown for 14 days before use. Approximately 10 seedlings per epiRIL were harvested (approx. 100 mg fresh weight) and DNA extracted using the Qiagen DNeasy Plant Mini Kit following the manufacturer’s instructions (Qiagen N.V., Hilden, Germany). Before library preparation, DNA was fragmented to an average size of 350 bp using 24 cycles of 30 s with a Bioruptor Diagenode sonication device. The genomic DNA sequencing libraries were prepared using the TruSeq DNA PCR-Free LT Library Prep Kit following the manufacturer’s instructions (Illumina, San Diego, Calif.), starting from 1.1 μg of fragmented DNA. Libraries were validated using High Sensitivity D1000 screentape on a 2200 Tapestation instrument (Agilent technologies, Santa Clara, CA) and a LightCycler 480 Instrument II using the LightCycler 480 SYBR Green I Master mix (Roche, Basel, Switzerland).

### DNA sequencing, reads mapping and peak call

The DNA libraries were sequenced with 2 × 76-bp paired-end reads with a minimum of 10× coverage (24× on average) on an Illumina NextSeq 500 using the High-Output Flow Cell configuration. The raw reads were trimmed using Trimmomatic to remove adapter sequences. Reads with an averaged value of at least 15 in a 4-nt window were trimmed from both ends. After trimming, reads pairs with at least one mate shorter that 36 bp were discarded. The remaining sequences (on average 95% of raw reads) were aligned with bowtie2 (Langmead and Salzberg, 2012) with the –no mixed and –non deterministic option, against the reference TAIR10 Arabidopsis genome (www.arabidopsis.org). Metrics of the sequencing analysis are displayed in Supplementary Table 8. Genome coverage was calculated for each genome position using genomecov (bedtools v2.26.0) and normalized to the Arabidopsis genome and library size. Arabidopsis genomic regions with a coverage of more than 2.5-fold or less than 0.2-fold of the average genome coverage in the Col-0 wild-type control were masked to avoid calling peaks in regions with high copy number or insufficient coverage (e.g. plastids DNA, ribosomal repeats, microsatellites). Peak call was done for each epiRIL mapped bam file in comparison to the Col-0 control condition with macs2 (https://github.com/taoliu/MACS/), setting the fragment size to 75 and extended size to 400 bp, with the options –B, -SPMR and -no model and Columbia-0 as control. The Log based background subtraction was performed with macs2 bdgcmp with “logLR” as model and a *P*-value threshold of 0.00001.

The peak lists obtained from all epiRILs were merged and peaks closer than 5 kb were joined together in a gff file using R (https://www.R-project.org/). Then, HTseq (Anders et al., 2015) was used to count reads at each peak for all conditions analysed and the raw count was normalized to peak length and library size to obtain FPKM values. Fold change (FC) of DNA coverage was calculated for each peak by the ratio between FPKM values in the tested condition and the Col-0 wild-type control. Peaks overlapping TEs (TAIR10 annotation) were further filtered to contain only regions with more than 1.8-fold change difference in at least one epiRIL.

### Detection of TE insertions

New putative TE integrations were detected in each epiRIL with the Transposon Insertion Finder (TIF) algorithm (Nakagome et al., 2014), using the first and last 17 bp of each TE tested and the Arabidopsis TAIR10 genome as reference. For each sample, the paired fastq files obtained from the genome sequencing were concatenated in a single file and used as input for TIF. The new putative insertion loci obtained from the analysis were filtered to exclude false positive calls in Col-0 sequenced samples.

### PCR and cloning

To confirm new transposon insertions, PCRs were carried out using a transposon-specific primer and a primer flanking the new insertion. Amplification of the entire locus with new Pack-CACTA1a integration was performed using two primers designed on the sequences flanking the new insertion. All PCR reactions were carried out using GoTaq enzyme (Promega, Madison, WI, USA), with extension time adjusted to the expected size of the fragment amplified, following the recommended manufacturer’s instructions. All PCR products were extracted from the gel using a Qiagen Gel Extraction Kit and eluted in 30 μl of water. The purified PCR products were ligated into a pGEM-T plasmid (Promega) following the manufacturer’s instructions and 2.5 μl of ligated sample used to transform E.coli DH5α cells (50 μl). Colony PCRs were performed starting from overnight grown *E. coli* colonies using the universal M13 forward and reverse primers. The amplified products were SANGER sequenced by Sigma-Aldrich (Merck, Darmstadt, Germany). Primer sequences are listed in Supplementary Table 9.

### RT PCR

Total RNA was extracted from 150 mg of fresh leaf tissue or green siliques using the Trizol (Invitrogen) method according to the manufacturer’s instructions and resuspended in 50 μl of water. Total RNA (10 μl) was treated with RQ1 DNase (Promega) and reverse transcribed using Superscript II (Invitrogen) following the manufacturer’s instructions. RT-PCR was carried out using specific primers designed on exons of the Pack-CACTA1a transposon and the gene target of integration (Supplementary Table 9).

### BLAST search

BLAST analyses were carried out using the NCBI web interface (https://blast.ncbi.nlm.nih.gov/Blast.cgi) or the locally installed BLAST+ v2.2.25 (Camacho et al., 2009), depending on the analysis. When the local version was used, blastn was run with the options–evalue 1e-5 and -max_target_seqs 100 where not specified otherwise.

### Pack-CACTA analysis

Additional Pack-CACTA elements were detected in the Arabidopsis genome by blasting the 26-bp core TIR sequences (Figure 1C) of the active Pack-CACTA1a element to the Arabidopsis reference genome (TAIR10). The blastn (BLAST+v2.2.25) was run with the options “-task blastn-short” and “-word_size 7”. The retrieved 18 matching sequences were imported in Geneious v7.1.7 (Biomatters) and manually checked to remove sequence derived from autonomous CACTA elements.

In order to identify Pack-CACTA elements in the genome of Ler-0 accession, the entire Pack-CACTA1a sequence was blasted on the PacBio Ler-0 assembly (http://www.pacb.com/). The sequences of the matching hits were retrieved from the assembly and aligned with Geneious v7.1.7 (Biomatters), using the MUSCLE algorithm and default parameters. Consensus trees were obtained from the alignment using the Temura-Nei genetic distance and Bootstrap resampling of 1000 replications. TIRs and TSDs were manually annotated.

### Rice Pack-MULE analysis

The annotation of rice Pack-MULEs was obtained by Jiang et al. (Jiang et al., 2011) and the sequence of each element was retrieved from the rice genome (http://rice.plantbiology.msu.edu/, release 6.0) using R. The sequences obtained were blasted against the same reference genome with blastn (BLAST+ v2.2.25) and the options-evalue 1e-5 and -outfmt 6. The list of 1,497,458 blast hits was filtered by alignment length > 500 and evalue < 1e-5, selecting only blast hits with “query_start < 20” or “query_end > (query_length – 20)” in order to select only results including at least one of the TIR sequences. The list was further filtered by excluding any sequence overlapping the original set of Pack-MULEs previously described (Jiang et al 2011). Remaining overlapping sequences were merged to generate the final list of 2,151 partial pack-MULE elements (Supplementary Data 8).

### Availability of data and materials

Sequencing data have been deposited in Gene Expression Omnibus under the accession number (sequencing data are under submission process). All the other data generated or analysed during this study are included in this published article or available under request.

## Author contributions

MC and JP designed the experiments; TJ prepared nucleic acid libraries; MC, TJ, and EC performed validation of new transposon insertions and their arrangement, MC performed all other experiments and analysis of the genomic data; MC and JP wrote the paper.

## Supplementary Figure legends

**Supplementary Figure 1.**
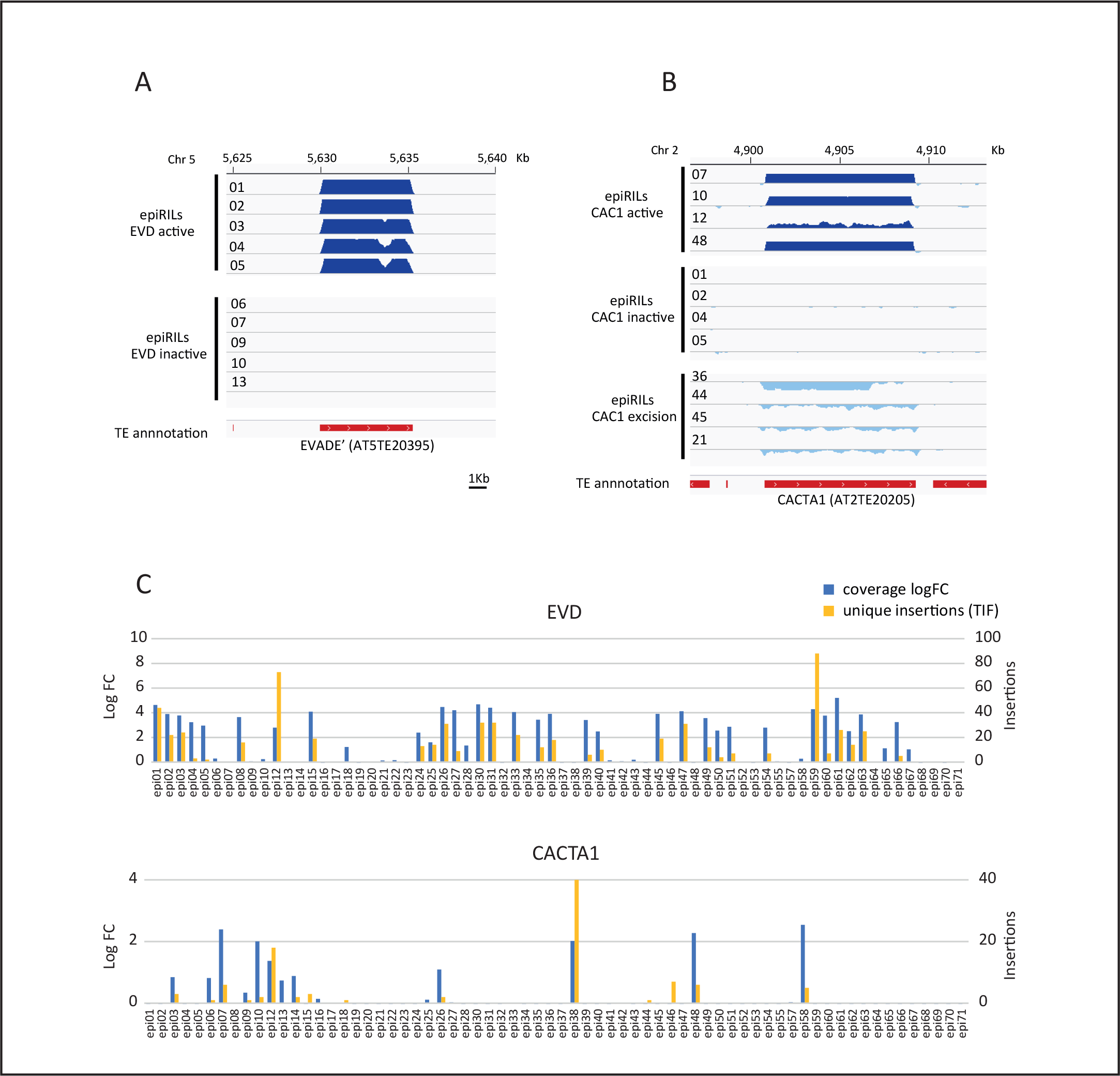
A) Copy number changes in epiRILs of EVADE’ retrotransposon. EpiRIL numbers are indicated at the left of each track. Log-transformed coverage difference of mapped reads at a locus between each of the indicated epiRIL and the reference Col-0 control. In each track, the relative coverage is plotted in intervals of 0 to 1. The track below indicates the TAIR10 annotation. B) Copy number changes in epiRILs of DNA transposon CACTA. EpiRIL numbers indicated at the left of each track. Log-transformed coverage difference of mapped reads at a CACTA1 locus between each of the indicated epiRILs and the reference Col-0 control. In each track, the relative coverage is plotted in intervals of −1 to 1. The decrease in coverage at some epiRILs observed is consistent with excision events occurring for DNA transposons. The track below indicates the TAIR10 annotation. C) Bar plot illustrating fold change (FC) in copy number of EVADE’ (upper plot) and CACTA1 (lower plot) transposons in each epiRIL line compared to Col-0. Blue bars represent DNA reads coverage and red bars represent number of novel “split reads”.

**Supplementary Figure 2.**
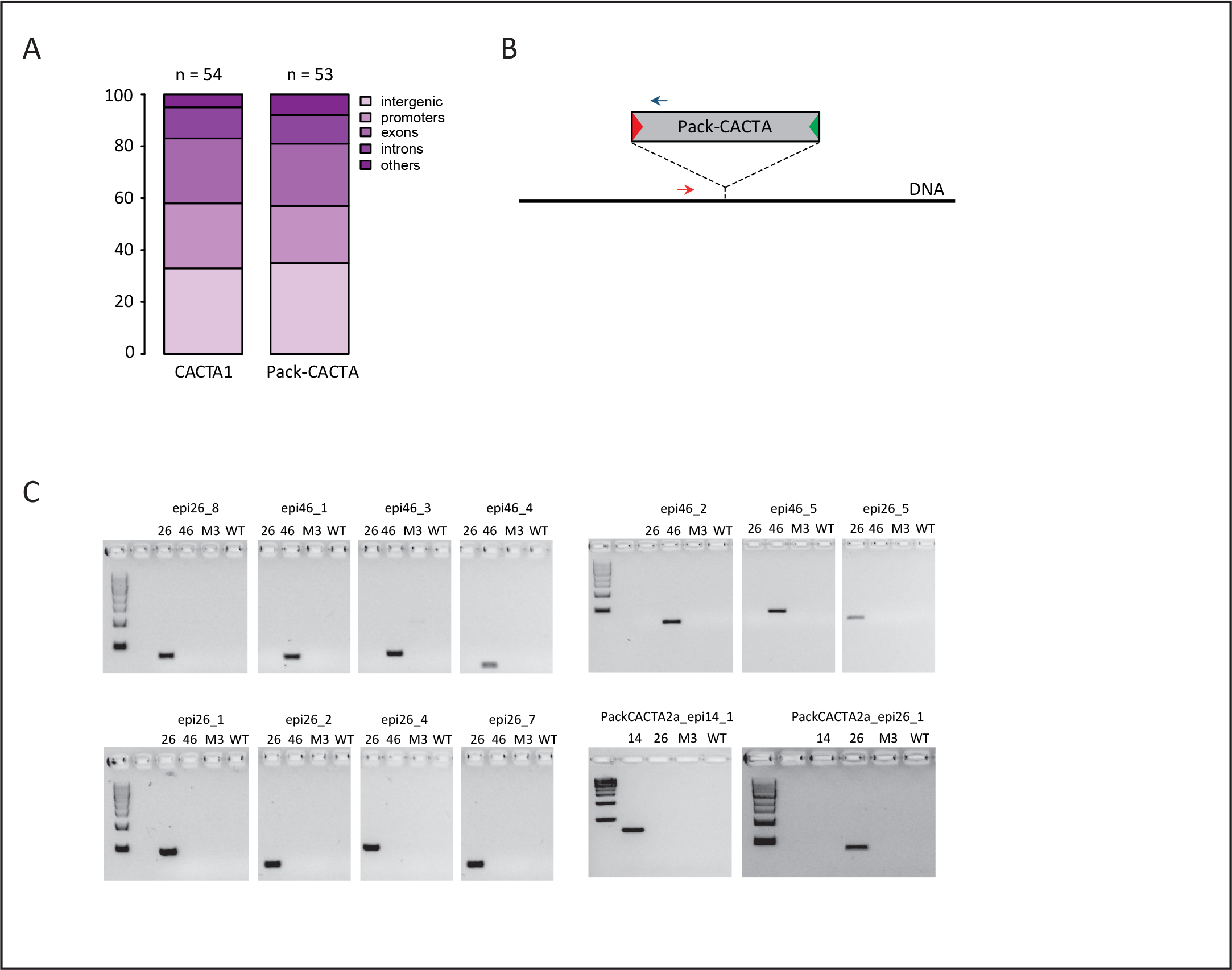
A) Chromosomal distribution of new insertions of CACTA1 and Pack-CACTA elements in Arabidopsis EpiRILs. Promoters were defined as stretches of maximum 1 kb DNA upstream of transcription starts of genes. B) Location of primers used for PCR validation of Pack-CACTA new insertions. C) Validation of new Pack-CACTA insertions in epi14 (14) epi26 (26) and epi46 (46). DNA from the original parental *met1-3* (M3) and Col-0 wild type (WT) are shown as negative controls.

**Supplementary Figure 3.**
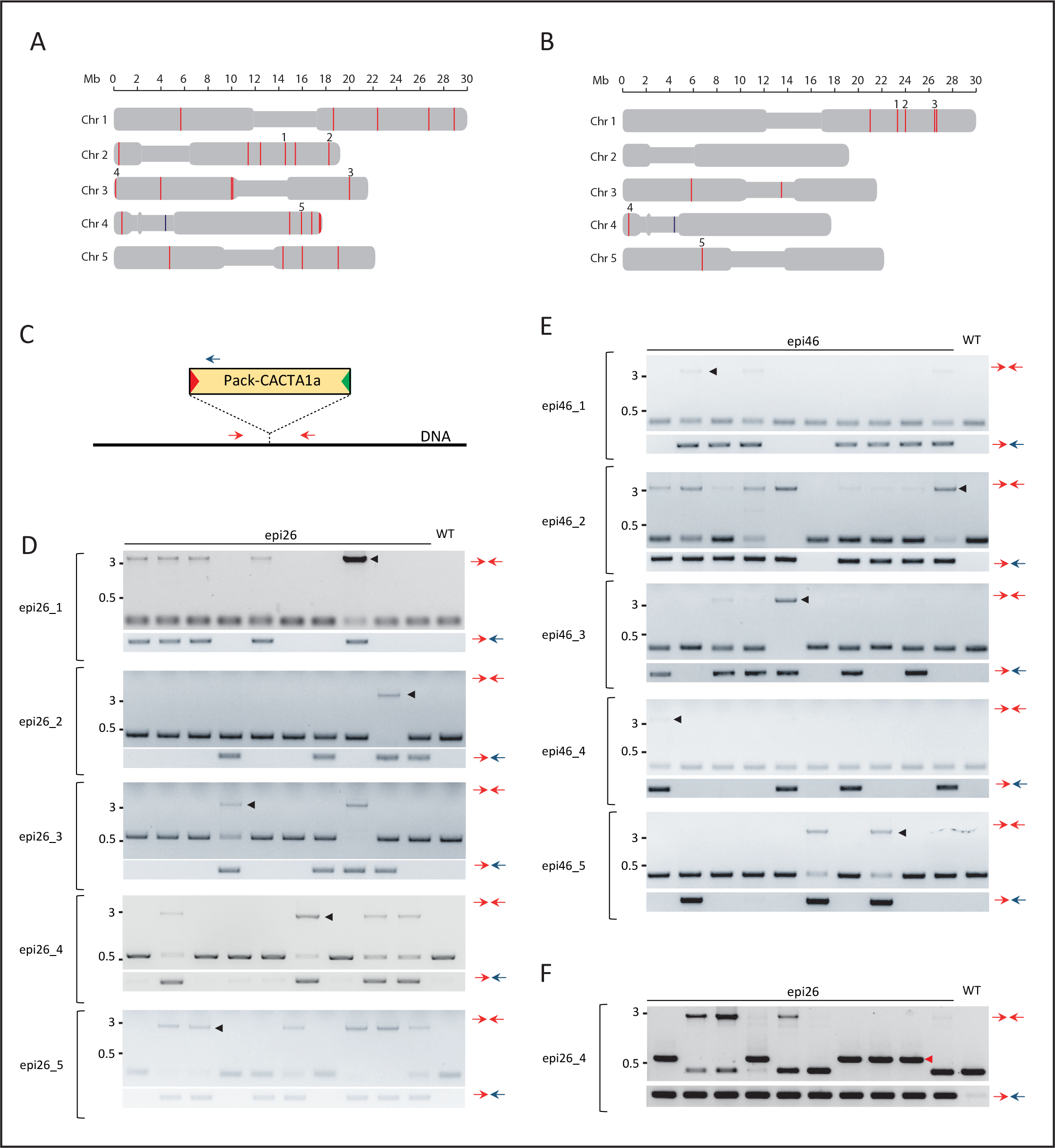
A) The positions of the loci amplified and cloned are indicated with numbers. The original Pack-CACTA1a copy on chromosome 4 is marked by a blue line. B) Distribution of new Pack-CACTA1a insertions in the epi46 genome (marked as in A). C) Strategy for PCR amplification of loci with a new Pack-CACTA1a integration. Locus-specific primers (red arrows) were designed for each new insertion studied. D) Amplification of five Pack-CACTA1a insertions in epi26 single plants. The primer combination used is represented by arrows on the right (for explanation see panel C). The black triangles mark the sequenced fragments confirming the presence of novel insertions of full-length Pack-CACTA1a. E) Amplification of five Pack-CACTA1a insertions in epi46 single plants (marking as in panel D). F) Amplification of the locus 4 in additional epi26 plants with a Pack-CACTA1a insertion, showing the presence of partially deleted transposons. A red triangular arrow indicates the sequenced Pack-CACTA1a deleted fragment. Description as in D.

**Supplementary Figure 4.**
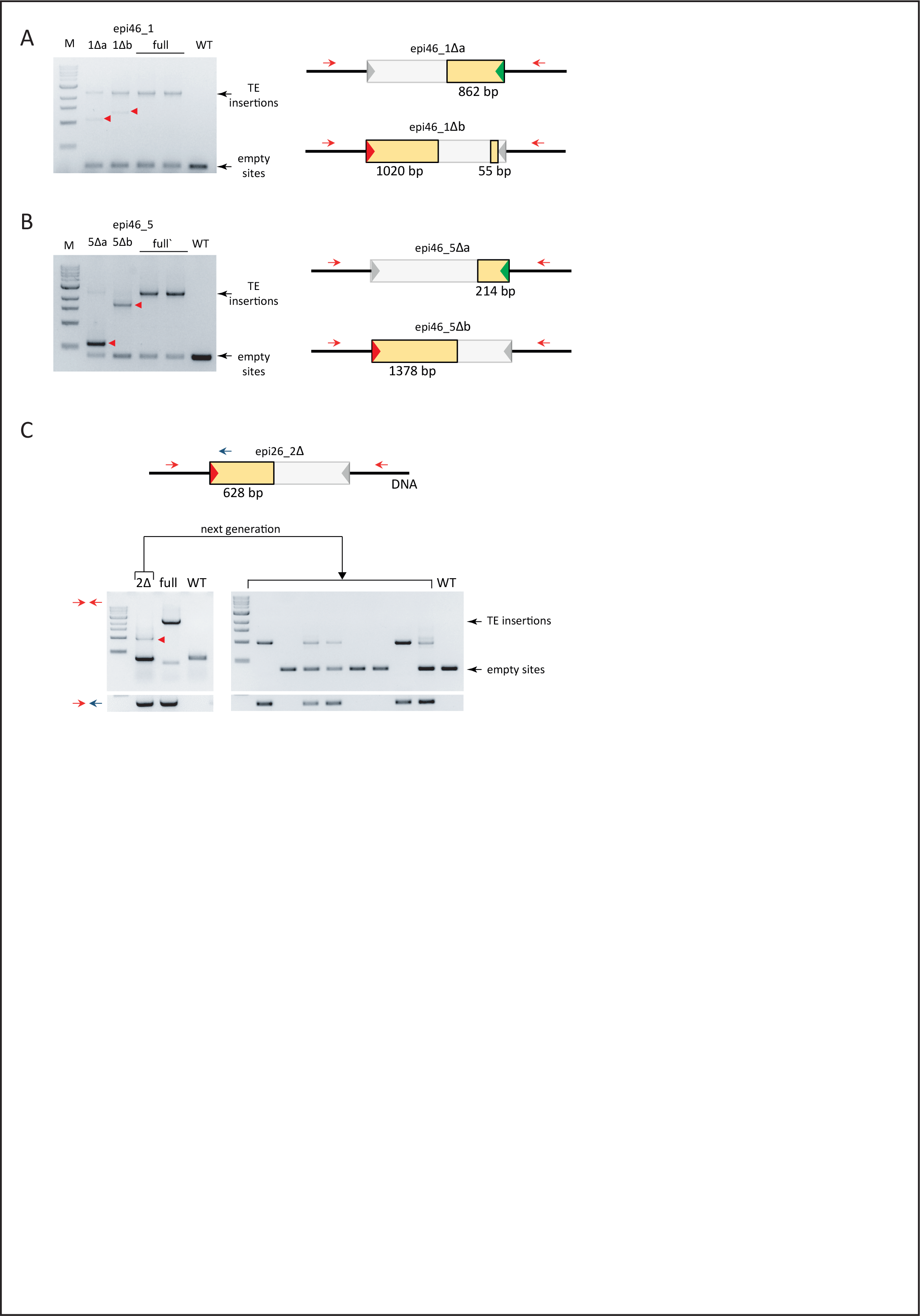
Formation of Pack-CACTA1a deletion derivatives in loci epi46_1 (A) and epi46_5 (B). PCR amplification products with deletion derivatives as marked by red triangles (left panels) and their schematic representations according to the obtained sequences (pictures on the right). Primers used are marked as red arrows. The ladder is 1 kb NEB (line M). Line WT represents Col-0. C) Formation and transgenerational transmission of a Pack-CACTA1a deletion derivative in locus epi26_2. The primers used are displayed as red and blue arrows. Red triangular arrows indicate the sequenced Pack-CACTA1a deletion derivative. Other markings as in A and B.

**Supplementary Figure 5.**
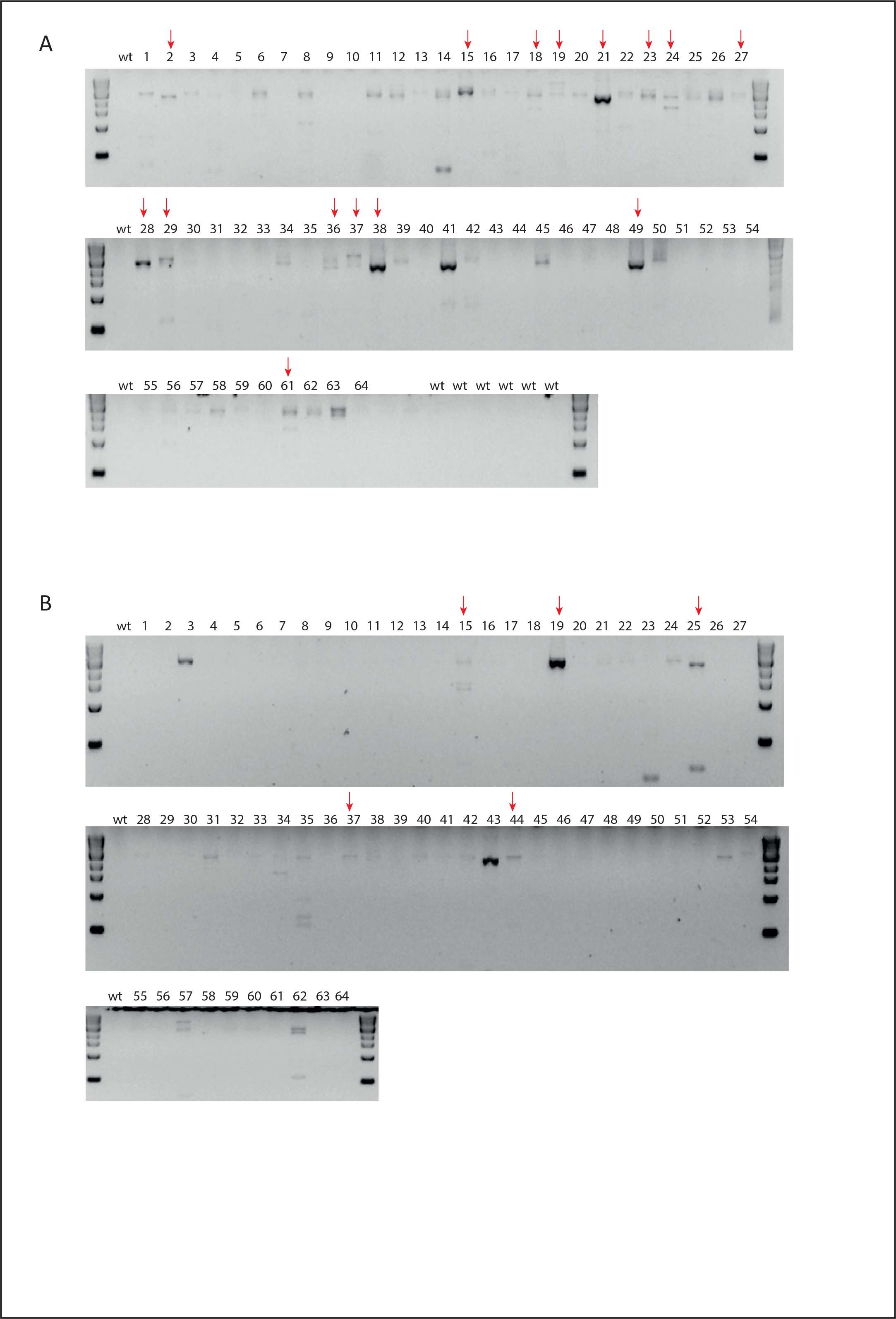
Results of PCR-based screening for closely spaced tandem insertions of CACTA1a (the strategy and primers as in Figure 4B). (A) Gel-separated PCR products representing 64 plants of epi26. (B) PCR products representing 64 plants of epi46. Red arrows point to samples subjected to sequencing. The upper gel in A represents a repetition of samples displayed in Figure 4B (note the much shorter separation of the PCR products in the gel presented here).

**Supplementary Figure 6.**
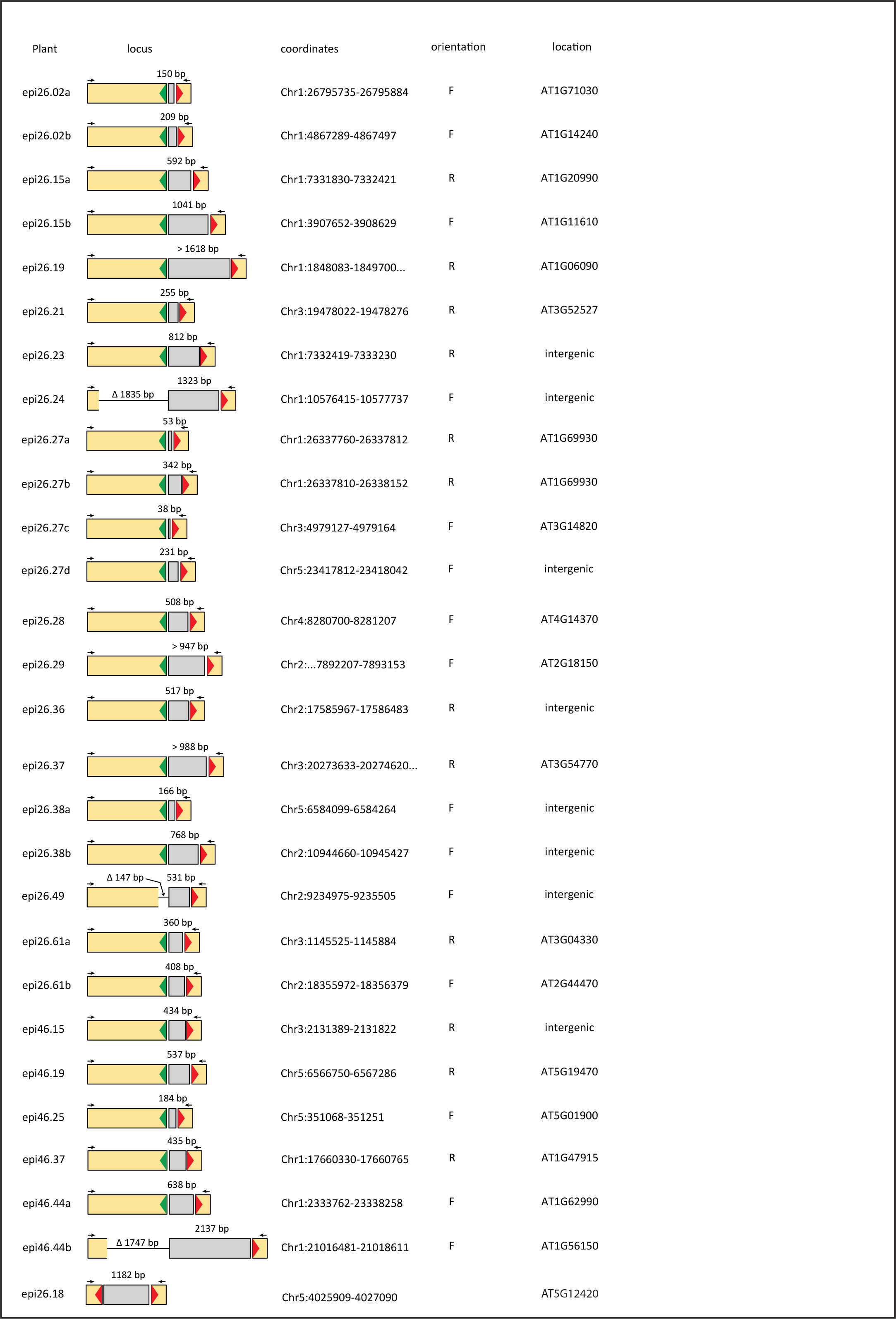
Schematic maps of 28 DNA sequences obtained from the 20 PCR reactions marked in Supplementary Figure 5. Red and green triangles symbolize 5’ and 3’ TIR, respectively. The positions of primers used for amplification are shown as black arrows and explained in Figure 4B. Closely spaced pack-CACTA1a insertions are predominantly arranged as direct repeats (27 examples of head-to-tail and only 1 head-to-head, epi26.18). Arabidopsis genomic DNA is displayed in grey and its coordinates orientation, and annotation are reported on the right.

**Supplementary Figure 7.**
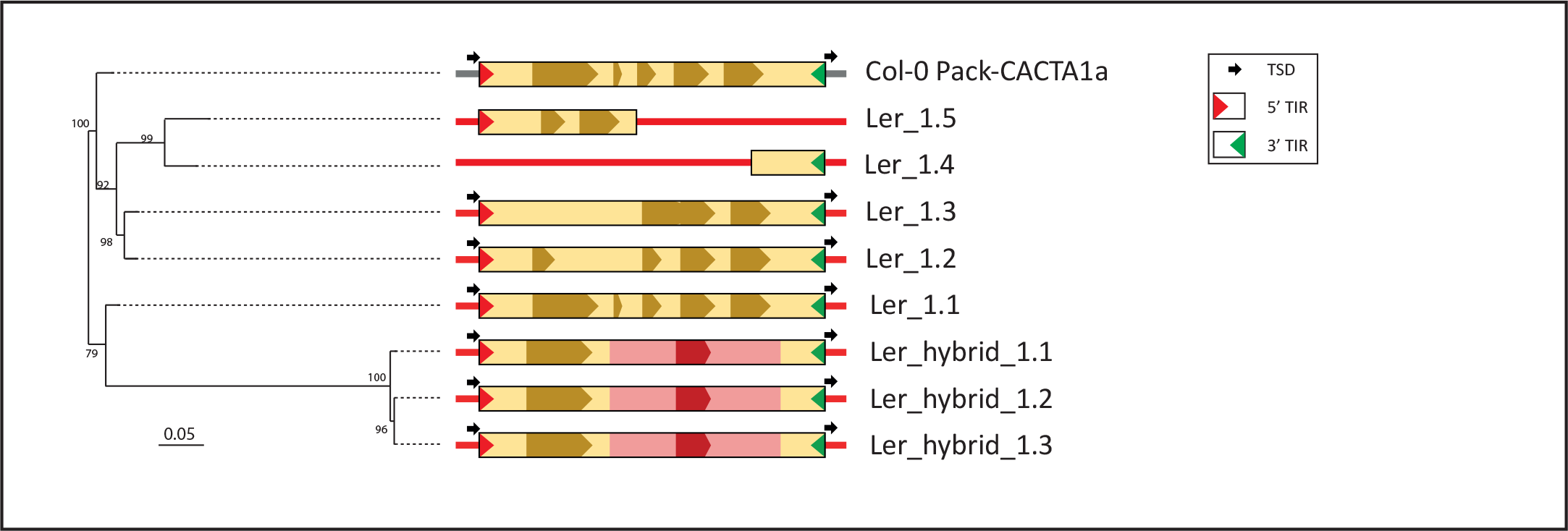
Structure of Pack-CACTA family in the Ler-0 accession of Arabidopsis. Exons are represented by darker hue. The phylogenetic tree was obtained using the alignment of full-length sequences and the numbers at each node indicate bootstrap support values of 1000 replications. Other symbols as in Figure 1B.

